## Supplementary material for "Computing 3D chromatin configurations from contact probability maps by Inverse Brownian Dynamics": Derivation of the Inverse Brownian Dynamics Method

### 1 Inverse Brownian dynamics

This section discusses the implementation of the Inverse Brownian Dynamics (IBD) method in the context of chromatin. The phase-space variable ($\Gamma$) for the coarse-grained bead-spring chain used to represent chromatin is the set of bead position vectors $\boldsymbol{r}_\mu(\mu = 1, 2, ..., N)$ and the Hamiltonian $\mathcal{H}(\Gamma, \{\epsilon_{\mu\nu}\}) = U^{\text{s}*} + U^{\text{SDK}*}$, where $U^{\text{s}*} = \sum\limits_{\mu=1}^{N-1} U_\mu^{\text{s}*}$ and $U^{\text{SDK}*} = \sum\limits_{\mu,\nu=1}^{N} U_{\mu\nu}^{\text{SDK}*}$. Since we are interested in bead-pairs, we construct a single index to represent any particular bead-pair. For instance, the expression

$$m = \frac{1}{2}[\mu(\mu - 1)] + [\nu - (\mu - 1)] \tag{1}$$

converts any bead pair $(\mu, \nu)$ to a single index $m$. Here $\mu$ varies from 2 to $N$, and $\nu$ varies from 1 to $(\mu - 1)$ for a matrix of size $N$. In terms of the single index, the average contact probability $p_m$ of the bead-pair $m$ is given by

$$p_m = \langle \hat{p}_m \rangle = \frac{1}{Z} \int d\Gamma\, \hat{p}_m \exp(-\beta\mathcal{H}) \tag{2}$$

Here, the partition function $Z = \int d\Gamma \exp(-\beta\mathcal{H})$, and $\hat{p}_m$ is an indicator function which indicates when contact occurs between the bead pair represented by index $m$. $\hat{p}_m$ is 1 if the distance between the beads is less than the cut-off distance of the indicator function, $r_\text{p}^*$, and 0 otherwise. For this work $r_\text{p}^* = r_\text{c}^*$, the cut-off distance of the SDK potential. We intend to target the experimentally obtained contact probability $p_m^{\text{ref}}$ by adjusting the well-depth of SDK attractive interactions $\epsilon_m$. The Taylor series expansion of $\langle \hat{p}_m \rangle$ about the interaction strength $\epsilon_m$ after neglecting higher order terms is

$$\langle \hat{p}_m \rangle(\epsilon_m + \Delta\epsilon_m) = \langle \hat{p}_m \rangle\, (\epsilon_m) + \sum_n \chi_{mn}\, \Delta\epsilon_n \tag{3}$$

where $\Delta\epsilon_m$ is the change in the interaction strength, and the susceptibility matrix

$$\chi_{mn} = \frac{\partial \langle \hat{p}_m \rangle}{\partial \epsilon_n} = \frac{\partial}{\partial \epsilon_n} \left[ \frac{1}{Z} \int d\Gamma \hat{p}_m \exp(-\beta\mathcal{H}) \right] \tag{4}$$

Simplifying further

$$\chi_{mn} = \frac{1}{Z}\left[\int \hat{p}_m \frac{\partial}{\partial \epsilon_n}\left(\exp(-\beta\mathcal{H})d\Gamma\right)\right] - \left[\frac{1}{Z}\left(\frac{1}{Z}\int \hat{p}_m \exp(-\beta\mathcal{H})d\Gamma\right)\frac{\partial Z}{\partial \epsilon_m}\right] \tag{5}$$

or

$$\chi_{mn} = \frac{\beta}{Z}\left[\int \hat{p}_m \exp(-\beta\mathcal{H})\frac{\partial(-\mathcal{H})}{\partial \epsilon_n}d\Gamma\right] - \beta\left[\langle \hat{p}_m\rangle \frac{1}{Z}\int \exp(-\beta\mathcal{H})\frac{\partial(-\mathcal{H})}{\partial \epsilon_n}d\Gamma\right], \tag{6}$$

Defining the quantity $b_n$ by

$$b_n = -\frac{\partial \mathcal{H}}{\partial \epsilon_n} \tag{7}$$

and using the expression for SDK potential, since only $U_n^{\mathrm{SDK}*}$ depends on $\epsilon_n$, one can show

$$b_n = \begin{cases} 1 & r_n^* \leq 2^{\frac{1}{6}}\sigma^* \\ \frac{1}{2}\left[1-\cos(\alpha\, r_n^{*2}+\beta)\right] & 2^{\frac{1}{6}}\sigma^* \leq r_n^* \leq r_{\mathrm{c}}^* \\ 0 & r_n^* \geq r_{\mathrm{c}}^* \end{cases} \tag{8}$$

which leads to

$$\chi_{mn} = \beta\left[\frac{1}{Z}\int \hat{p}_m b_n \exp(-\beta\mathcal{H})d\Gamma - \langle \hat{p}_m\rangle \frac{1}{Z}\int b_n \exp(-\beta\mathcal{H})d\Gamma\right] = \beta\left[\langle \hat{p}_m b_n\rangle - \langle \hat{p}_m\rangle\langle b_n\rangle\right] \tag{9}$$

Replacing the left hand side of Eq. 3 with the target contact probability $p_m^{\mathrm{ref}}$ obtained from experiment, we get

$$p_m^{\mathrm{ref}} - \langle \hat{p}_m\rangle = \sum_n \chi_{mn}\,\Delta\epsilon_n \tag{10}$$

Equation 10 can be solved for any particular iteration step as

$$\epsilon_n^{(i+1)} = \epsilon_n^{(i)} + \lambda \sum_m C_{nm}^{(i)}\left(p^{\mathrm{ref}} - \langle \hat{p}_m\rangle^{(i)}\right) \tag{11}$$

where the matrix $\mathsf{C}$ is the *pseudo-inverse* of the matrix $\chi$, superscript $i$ represents the iteration number, $\lambda$ denotes the damping factor with $0 < \lambda < 1$, and $\epsilon_n^{(i+1)}$ is the well-depth of the SDK attractive interaction for the next iteration step. Since the susceptibility matrix $\chi$ is often singular (with the rank of the matrix being smaller than its size), it is necessary to define the matrix $\mathsf{C}$ as a *pseudo-inverse* and determine it by using *singular-value decomposition* (SVD). By carrying out SVD, the susceptibility matrix is decomposed into three square matrices $\mathsf{U}, \mathsf{S}$ and $\mathsf{V}$ such that $\chi = \mathsf{U}\mathsf{S}\mathsf{V}^{\mathrm{T}}$ where $\mathsf{U}$ and $\mathsf{V}$ are orthogonal and $\mathsf{S}$ is a diagonal matrix. The *pseudo-inverse* of the susceptibility matrix via SVD can then be written as

$$\mathsf{C} = \chi^{-1} = \mathsf{V} \cdot [\text{diag}(1/S_n)] \cdot \mathsf{U}^T \tag{12}$$

Note that in all the cases considered here, $\chi, \mathsf{S}, \mathsf{U}$ and $\mathsf{V}$ are square matrices since there is a one-to-one correspondence between contact probability and interaction strength. As per the well-established procedure for finding the *pseudo-inverse*, when $S_n < 10^{-4}$, $1/S_n$ is replaced by 0. In other words, some values in the $\mathsf{S}$ matrix whose inverse would lead to problems, are ignored.

An example of the $\mathsf{S}$ matrix obtained in the case of the prototype scenario where we have considered a bead-spring chain with 45 beads is given here as illustration. The matrix is of size $N(N-1)/2 \times N(N-1)/2 = 990 \times 990$, which represent the number of interacting bead-pairs in the chain. Fig. S1 represents the matrix at an intermediate stage in the iteration process. It can been seen that $\mathsf{S}$ has diagonal elements whose magnitude is of order $10^{-4}$ or less (top right in Fig S1).

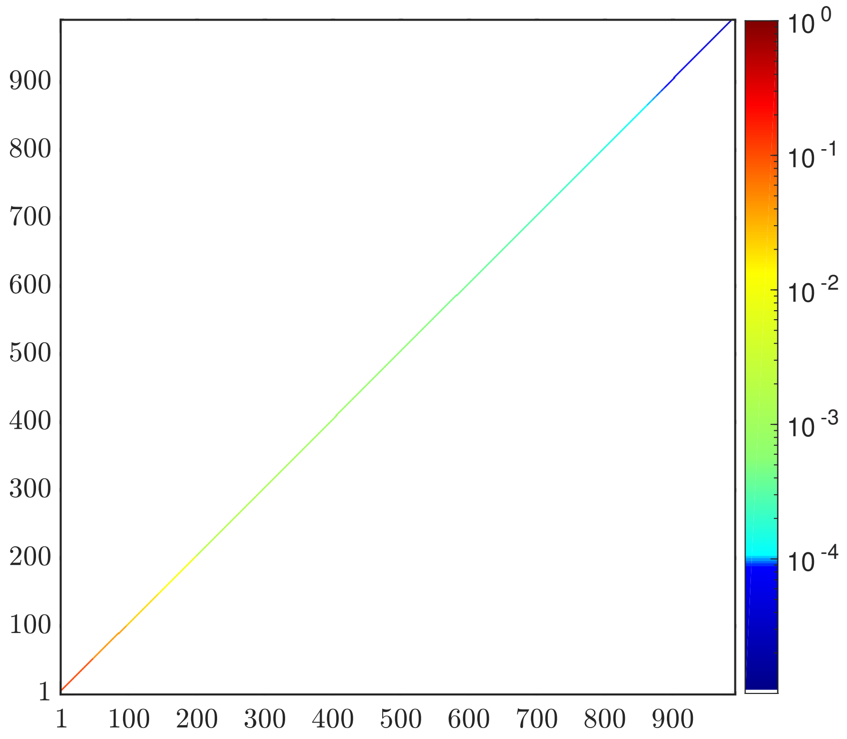


Figure S1: Representation of the matrix $\mathsf{S}$. The top right corner along the diagonal represents diagonal elements of the matrix, $S_n$, which are of order $10^{-4}$ or less.
